# Geometric optics organizes organelle interactions and positioning

**DOI:** 10.64898/2026.04.28.721423

**Authors:** Reza Farhadifar, Gunar Fabig, Hai-Yin Wu, Yuan-Nan Young, Justus Vogel, Daniel J. Needleman, Thomas Müller-Reichert, Michael J. Shelley

**Affiliations:** Center for Computational Biology, Flatiron Institute; New York, NY, USA; Experimental Center, Faculty of Medicine Carl Gustav Carus, Technische Universität Dresden; Dresden, Germany; Department of Molecular and Cellular Biology, Harvard University; Cambridge, MA, USA; Department of Mathematical Sciences, New Jersey Institute of Technology; Newark, NJ, USA; Department of Visual and Data-Centric Computing, Zuse Institute; Berlin, Germany; Core Facility Cellular Imaging, Faculty of Medicine Carl Gustav Carus, Technische Universität Dresden; Dresden, Germany; Courant Institute, New York University; New York, NY, USA

## Abstract

Microtubule asters are central to positioning organelles and organizing intracellular architectures. We show that the multi-stage choreography of microtubule asters which positions pronuclei during the first cell division in *Caenorhabditis elegans*, is governed by a cellular analog of geometric optics. Large-scale electron tomography reveals that astral microtubules reach cortical and pronuclear surfaces largely along line-of-sight trajectories, leaving complementary regions shadowed and inaccessible. Laser ablation identifies surface-anchored motors pulling on microtubules as the dominant drivers of motion. We develop a biophysical model incorporating dynamic terminator curves that separate microtubule-accessible and -inaccessible surfaces, and quantitatively recapitulate aster separation, pronuclear migration, centering, and rotation in control and genetically perturbed embryos. More broadly, robust positioning emerges from a feedback loop wherein intracellular geometry gates microtubule access, thus sculpting the distribution and magnitude of pulling forces, which in turn reshape that geometry.

## Main Text

Proper positioning of a cell’s contents is essential to successful cell division and the development of an organism. One prominent example is the precise migration of pronuclei, structures that carry maternal and paternal genetic information, in early embryos of the nematode *Caenorhabditis elegans* (*1*). Following fertilization, the two asters (each composed of a centrosome and the microtubule array emanating from it) migrate to opposite positions on the male pronucleus, guiding fusion with the female pronucleus. The resulting pronuclear complex then moves to the cell’s center and aligns along the embryonic axis. This prototype dynamics ensures proper division. Underlying these coordinated motions are molecular motors, anchored to different surfaces within the cell, that generate forces on microtubules. However, because pronuclei can both carry motors, move relative to other structures, and possibly block microtubule access to surrounding surfaces, understanding these dynamics requires considering not only forces, but also an evolving three-dimensional geometry.

A recurring theme in many phenomena in chemistry, physics, and biology is of interactions between extended objects that are shaped by which parts of their surfaces are mutually accessible. For example, in directional thin-film deposition, atoms or molecules arrive along nearly straight trajectories, so only surface regions with line-of-sight to the incoming flux accumulate material, while shadowed regions remain uncoated, making surface growth strongly geometry-dependent (*2*). On the Earth’s surface, the distribution of heating is likewise determined by which regions lie on the day side and which lie in darkness, and the moving day-night boundary continually reshapes the pattern of energy input that drives atmospheric circulation (*3*). In phototropic plant growth, stems reorient toward a light source because photoreceptor activation is confined to the illuminated side, causing auxin to accumulate on the shaded side and drive greater cell elongation there; the moving boundary between illuminated and shaded tissue therefore sets where growth is enhanced and how the stem curves over time (*4*).

The common feature in these examples is that one can draw on a surface a boundary that separates regions that have direct line-of-sight to a given source from those that are shadowed. This boundary is known as a *terminator curve,* and its motion shifts surface elements from being accessible to inaccessible and *vice versa*. Here, we combine large-scale serial-section electron tomography, diffraction-limited femtosecond laser ablation, and rigorous computational modeling to show that a dynamic force balance, governed by moving terminator curves, coordinates aster separation and pronuclear migration in early *C. elegans* embryos.

### The geometry of aster engagement with pronuclei

How do microtubules from a nearby aster engage with a large, curved pronucleus? Geometrically, two scenarios are plausible. In one scenario, microtubules grow radially from the centrosome and remain essentially straight, contacting only those regions of the pronucleus that lie along their line of sight. In this case, the contact regions of microtubules with the pronucleus surface lie predominantly close to each aster, and regions in the geometric shadow remain inaccessible. In the other scenario, microtubules bend around the pronuclear envelope, effectively giving the asters access to almost the entire surface of the pronucleus. These two scenarios will give rise to fundamentally different conceptions of how forces are induced by motor activity onto asters.

To distinguish between these scenarios, we used large-scale electron tomography to visualize how astral microtubules are organized near the male pronucleus during its migration and when the female and male pronuclei meet (Fig. 1A-D) (*5*). We assembled six large-scale volumetric tomographic data sets spanning four key time points: when the two asters are tightly positioned between the male pronucleus and the posterior cortex (Fig. 1A and movie S1); when the asters have moved to opposing sides of the male pronucleus (Fig. 1B and movie S2); when the male and female pronuclei migrate towards one another (Fig. 1C and movie S3); and when the pronuclei meet and the asters are positioned in the groove between them (Fig. 1D and movie S4). By identifying centrosomes, tracing individual microtubules, and segmenting the pronuclear envelope, we obtained reconstructions that reveal the number, morphology, and spatial distribution of microtubules near the pronuclei. We use this data to investigate the geometry of aster-pronucleus engagement (Materials and Methods).

**Fig. 1.**
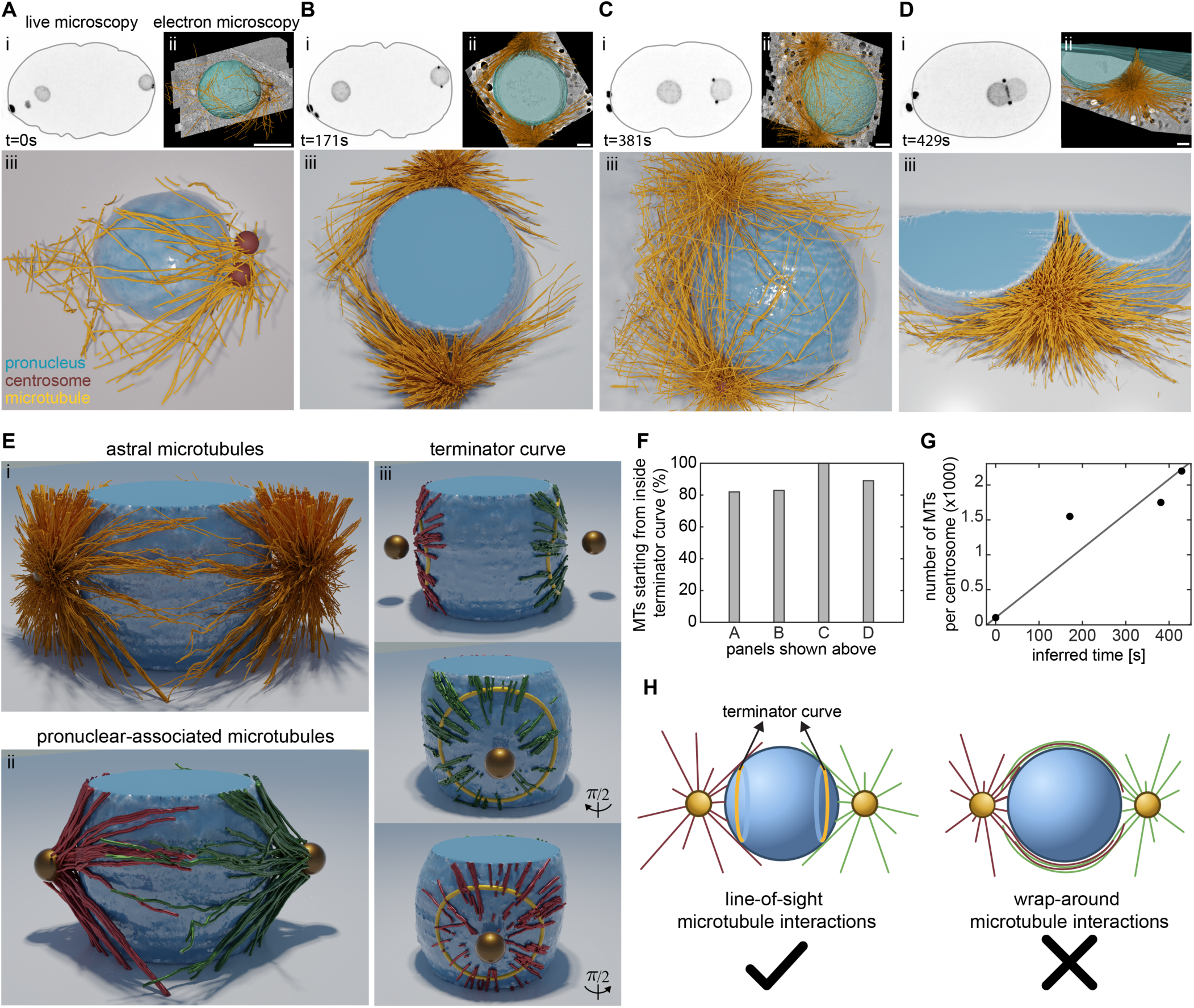
Microtubule organization during pronuclei migration. (**A**–**D**) (i) From live-cell microscopy, snapshots showing centrosomes and pronuclei at different stages of aster separation and pronuclear migration. The cell periphery is highlighted with a gray line. Scale bar, 10 μm. From electron tomography: (**A**–**D**) (ii) Overlay of segmented pronuclei (cyan), microtubules (orange), and centrioles (blue) with tomographic slices at the inferred time points in (i). Scale bar, 2 μm. (**A**–**D**) (iii) 3D rendering of pronuclear envelope (blue), centrosomes (red), and microtubules (orange) from the tomography reconstruction shown in (iii). (**E**) (i) 3D rendering of pronuclear envelope (blue), centrosomes (gold), and microtubules (orange) from the tomogram shown in **B**, shown from the side. (ii) Same tomogram and rendering angle as in (i), showing the subpopulation of microtubules that are 120 nm or closer to the pronucleus surface. (iii, top panel) Same tomogram and rendering angle as in (ii), showing only regions of microtubules that are 120 nm or closer to the pronuclear surface. Computed terminator curves are shown in yellow. The same rendered images from two different angles are also shown in (iii). (**F**) Fraction of microtubules first contacting the pronuclear surface inside the terminator curves for tomograms shown in (**A-D**). (**G**) Number of microtubules measured from tomograms as a function of time inferred from light microscopy. (**H**) Schematic of line-of-sight interactions vs wrap-around interactions of microtubules and the pronucleus.

We take the stage shown in Figure 1B, where the two asters are at opposing sides of the male pronucleus, as an example (viewed from a different angle in Fig. 1E). From all the traced astral microtubules (Fig. 1E-i), we first identified pronuclear-associated microtubules as those whose trajectories at any point approached the pronuclear envelope closer than 120 nm (Fig. 1E-ii). We then focused on the segments of these microtubules that are close to the pronuclear surface (distance is less than 120 nm), as shown in Figure 1E-iii. These segments form two compact caps facing their respective asters and are separated by broad regions with no microtubule associations. The outer edge of each cap is sharp: almost all contacts lie inside a well-defined boundary, with none wrapping around to the opposite side of the pronucleus. This pattern is consistent with the line-of-sight scenario and suggests that a geometric boundary on the pronuclear surface can be used to separate regions that are accessible and inaccessible to astral microtubules engaging with motors. Accordingly, we introduce the terminator curves which, for each aster, is defined by intersecting the pronuclear surface with a cone whose apex is at the centrosome and whose surface is tangent to the pronucleus. Overlaying the terminator curve thus defined on the tomographic reconstructions, we find that more than 80% of pronuclear-associated microtubules first contact the pronucleus surface inside the terminator curve (Fig. 1E-iii, 1F). For some microtubules, the entire contact region lies within the terminator curve, and for some, it extends slightly beyond. We observe similar line-of-sight interactions of asters with pronuclei for other tomographic reconstructions as well (Fig. 1F, Fig. S1-S3).

During the period of pronuclear migration, centrosomes are known to mature and increase in size (*6*), and this maturation is confirmed by light microscopy, which shows a rise in tubulin intensity around the centrosomes (*7*). Consistent with this, our tomograms show an increase in the number of astral microtubules over time: rising from ∼100 microtubules at early stages (Fig. 1A) to ∼2,200 by the time the pronuclei meet (Fig. 1D). By aligning the relative positions of asters and pronuclei in our tomograms with live-cell imaging, we can approximately time-order the reconstructions and infer that the number of microtubules increases by about 20-fold over ∼400 s (Fig. 1G).

Taken together, the electron tomography data show that a line-of-sight construction based on terminator curves effectively captures the geometry of the interaction between asters and pronuclei (Fig. 1H). For each aster, most pronuclear-associated microtubules first come close to the pronuclear surface inside that aster’s terminator curve, and only some contacts extend beyond this boundary. Except at the earliest time point when the two asters are closely apposed, the contact regions from the two asters remain largely separated on the pronuclear surface and do not intermingle. With this geometric framework in hand, we next probe the mechanics of pronuclei migration and centering in early *C. elegans* embryos.

### Cortical and pronuclear pulling forces act on asters to drive pronuclear migration

The two asters in the early *C. elegans* embryo are the main agents of force transduction and the dynamics studied here. Astral microtubules emanating from each aster contact the male and female pronuclei and the cell cortex, allowing forces generated on these surfaces to act through the asters and thereby move and orient the pronuclear complex. Our tomographic analysis shows that these contacts are strongly constrained by line-of-sight geometry: as asters and pronuclei move relative to one another, the regions of each surface that are accessible to astral microtubules, and thus able to exert force, are continually reshaped. This geometric view specifies where forces can be transmitted, but not how they are generated. It is unclear to what extent the microtubules which contact these surfaces are subject to pushing forces, exerted by microtubule polymerization or bending, or pulling forces, exerted by motors anchored to the surfaces (*8, 9*). To quantitatively examine how different forces contribute to pronuclear motion within this evolving geometric framework, we use a femtosecond stereotactic laser ablation system (FESLA) as a versatile and precise tool for dissecting when, where, and in what direction forces are exerted (*10–12*).

We first focused on the configuration in which the two asters have moved to opposing sides of the male pronucleus (Fig. 1B), and asked what forces the asters experience from the male pronucleus and from the cell cortex. In this geometry, microtubules between the aster and the male pronucleus could, in principle, either push the aster away from the pronucleus or pull it toward it, and cortical microtubules could likewise push or pull. To distinguish between these possibilities, we used FESLA to sever microtubules lying between the aster and the male pronucleus while performing fast 3D imaging before and after ablation, allowing us to directly probe forces on the aster based on its motion after ablation (Fig. 2A and movie S5). With ablation, this aster, which was initially near the male pronucleus, moved away from the pronucleus and towards the cell cortex, reaching a maximal distance of ∼8 μm in ∼10s before returning to the pronucleus (Fig. 2A-ii). Repeating this ablation experiment shows unambiguously that, at this time during pronuclear migration, the aster is being pulled by motors on both the cortex and the male pronuclear surface (Fig. 2B): cutting the microtubules yields an unbalanced cortical pull that drives the aster toward the cortex. Subsequent movement towards the pronucleus is consistent with the re-establishment of a balance between these two pulling forces that positions the aster near the pronucleus (Fig. 2B-iv).

**Fig. 2.**
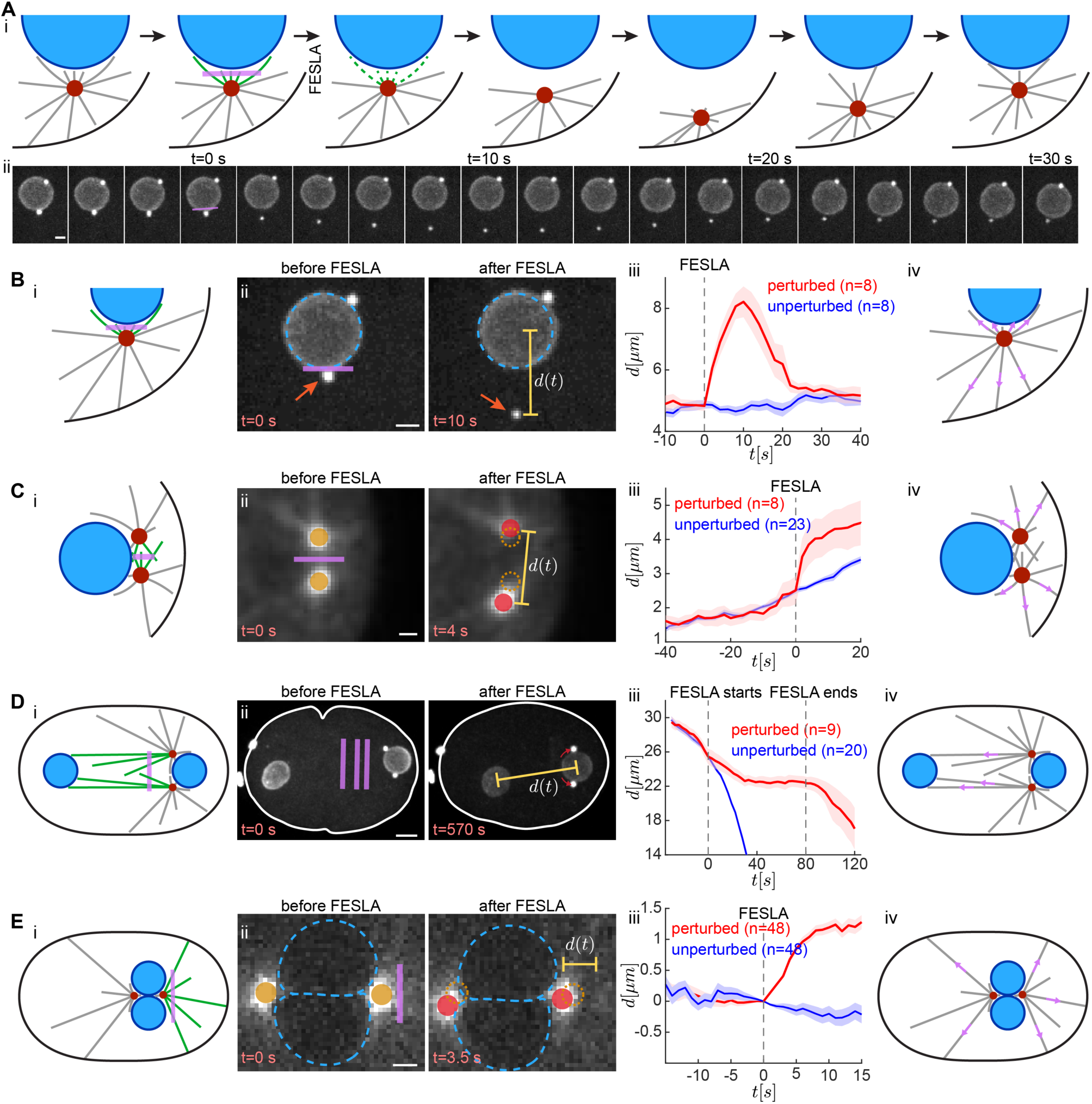
Pulling forces drive aster separation and pronuclear migration. (**A**) Severing of microtubules between the aster and male pronucleus using femtosecond stereotactic laser ablation (FESLA). The schematic in (i) shows severed microtubules (green), intact microtubules (gray), the centrosome (red), the pronucleus (blue), and the cell cortex (black). Snapshots before and after ablation are shown in (ii). Scale bar, 2 μm. (**B**–**E**) (i) Schematic of the ablation geometry, with severed microtubules (green), intact microtubules (gray), and ablation regions (purple). (**B**–**E**) Snapshots from before and after FESLA are shown in (ii). Note that in (D), multiple rounds of FESLA were performed. Scale bars: (B) 2 μm; (C) 1 μm; (D) 5 μm; (E) 2 μm. (**B**–**E**) (iii) Average distance over time for perturbed motion following FESLA (red line: mean; shaded area: standard error of the mean) and unperturbed controls (blue line: mean; shaded area: standard error of the mean). (**B**–**E**) (iv) Schematics showing the inferred direction of forces based on each corresponding FESLA experiment.

We next asked what forces the two asters exert on each other when they are initially close (Fig. 2C). When they are relatively close, the microtubules growing from each aster could, in principle, push the other aster away. In such a model, plus-end-directed motors such as kinesin-5 would generate anti-parallel sliding between overlapping microtubule arrays, producing a repulsive force between the two asters. If so, we anticipate that the ablation of the microtubules between the two asters slows down or stops their separation. To test this, we applied FESLA to sever the microtubules in between the two asters and instead observed that their separation was even faster than when unperturbed (Fig. 2C-i, ii, iii, and movie S6). This experiment argues that there is little such pushing force between asters (Fig. 2C-iv), consistent with previous studies showing that perturbation of *bmk-1* (a homolog of kinesin-5 in *C. elegans*) had little effect on aster separation in early *C. elegans* embryos (*13*).

So far, we have shown that motors on the cell cortex and on the male pronucleus pull on the asters. What forces does the female pronucleus exert on the asters? The onset of posterior-directed migration of the female pronucleus coincides with the positioning of the asters on opposing sides of the male pronucleus, placing the female pronucleus in direct line-of-sight of astral microtubules. We hypothesized that motors anchored to the female pronucleus pull on astral microtubules and move the female pronucleus toward the aster/male pronucleus complex. To test this, we repeatedly applied FESLA to sever microtubules between the two pronuclei when they were ∼25 μm apart. During ablation, posterior-directed migration of the female pronucleus ceased (Fig. 2D-i, ii, iii, and movie S7). Once we stopped ablation, migration resumed at a speed similar to the unperturbed embryo (Fig. 2D-iii). This experiment shows that the female pronucleus is also covered with motors and is being pulled upon by the asters (Fig. 2D-iv). Moreover, these pulling forces operate over distances of tens of microns, indicating that at least some astral microtubules grow to such lengths or that a network of connected microtubules spans this distance (*14, 15*).

Finally, we investigated the forces that center the joined aster/pronuclei complex. To do this, we used FESLA to cut astral microtubules that presumably connect the asters to the posterior cortex (Fig. 2E-i). Following ablation, the complex moved away from the cut region, indicating that pulling forces from the cortex contribute to centering at later stages (Fig. 2E-ii, iii, iv, and movie S8).

Overall, our ablation experiments argue that asters are pulled by motors distributed on the surfaces of the cell and the pronuclei. Presumably, the balance of these forces, perhaps with other forces such as cytoplasmic drag, maintains the asters near the male pronucleus in early stages, then drives their separation, orchestrates the migration of the male and female pronuclei, and then drives the positioning and orientation of the pronuclear complex. We argue that these forces depend strongly and nonlinearly on the evolving cellular geometry. This geometry comprises, among other things, the cellular and pronuclear shapes and sizes, and their positions relative to the force-bearing astral microtubules. We now turn to modeling concisely how this geometry evolves.

### A geometric optics model for pronuclear positioning

Our experiments support a physical picture for how asters interact with pronuclei and the cell cortex. Microtubules emanate from each aster in all directions, analogous to rays from a light source, and can engage motors anchored to the pronuclear envelope or the cell cortex as they probe the cytoplasm. The resulting motor-generated forces, balanced by cytoplasmic drag, act on both asters and pronuclei and drive their motion. Crucially, this force generation is geometrically controlled, being permitted only where microtubules have an unobstructed line of sight to a surface. For each aster, the boundary between a surface region reachable by microtubules from a region in shadow defines a terminator curve on that surface, directly analogous to terminator curves in geometric optics that separate illuminated and shadowed regions relative to a light source. As asters and pronuclei move, these terminator curves continuously reconfigure, thereby determining motor accessibility and the resulting forces and torques.

To capture the dynamics, we developed a geometric-optics based model (the GO-model) of aster and pronuclear motion. In the model we account for cell shape, two mobile and spherical pronuclei, and two mobile asters. We assume that all surfaces, the cell cortex and the pronuclei, are uniformly populated by motors, with *M_c_* motors on the cell cortex and *M_p_* motors on each of the pronuclei (Fig. 3A). Microtubules nucleate uniformly from the two asters at a rate *ṅ*(*t*) = 10 + *t*/2, grow outwards at their plus-ends with a speed *V_g_*, and undergo catastrophe at a rate *λ*. For simplicity, we assume all motors possess identical biophysical properties. The binding of a motor to microtubules is taken as explicitly stoichiometric: each motor is constrained to bind only one microtubule at a time. We have shown that this assumption, reflected by the structure of dynein motors, can provide stable positioning of the mitotic spindle in the presence of purely pulling forces, partly by inducing a competition between multiple asters seeking to occupy a motor (*10, 16*). When a motor at a surface position ***Y*** binds a microtubule originating from an aster at position ***X****_a_*, the motor exerts a pulling force 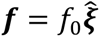, where *f*_0_ is the force magnitude and 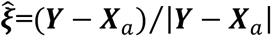 is the unit orientation vector from the aster to the motor.

**Fig. 3.**
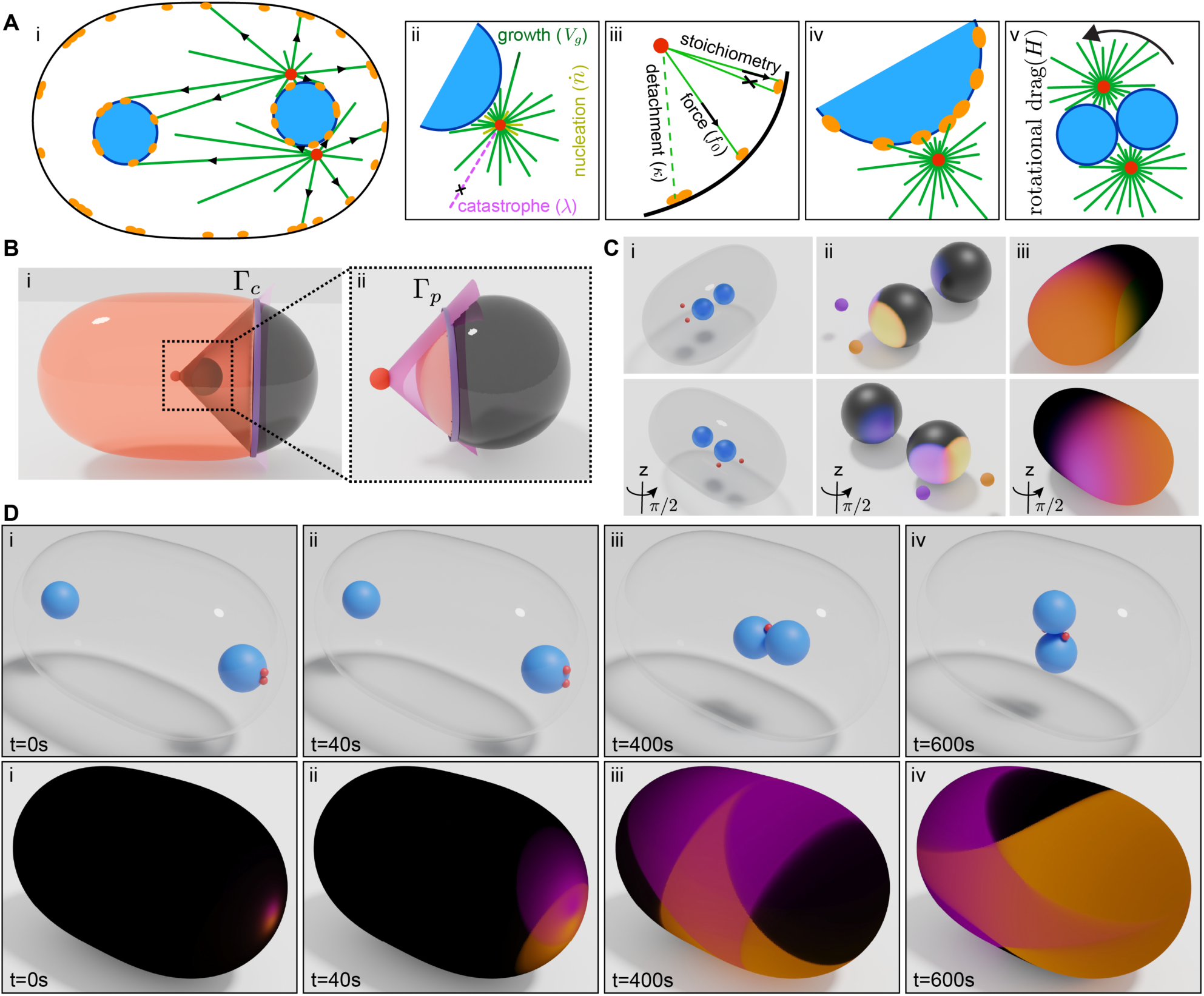
The GO-model of asters and pronuclei. (**A**) Schematic of the model showing centrosomes (red), microtubules (green), pronuclei (blue), and motors (orange). Black arrowheads show microtubules under pulling forces from motors. Key model elements are indicated: microtubule nucleation (rate *ṅ*), microtubule tip growth (speed *V_g_*), microtubule catastrophe (rate *λ*), stoichiometric attachment of microtubules to a motor, pulling force by a bound motor on a microtubule (*f*_0_), and detachment of microtubules from bound motors (rate *κ*). Motors are distributed uniformly over both the cell periphery and the pronuclear surfaces. Once the pronuclei meet, they join and can rotate with rotational drag coefficient *H*. (**B**) Schematic of terminator curves (Γ_*c*_ and Γ_*p*_) on the cell surface and the pronuclear surface, respectively, with the line-of-sight cone of motor engagement shown (purple). Orange regions indicate cell and pronuclear surfaces accessible to centrosomal microtubules, while dark gray labels inaccessible surfaces. (**C**) For the given arrangement of centrosomes and pronuclei shown in (i), two views are shown in (ii) and (iii) for the binding probability of motors on the pronuclear surface and cell surface, respectively. (**D**) From a full simulation of the GO-model, snapshots of the evolution of centrosomes and pronuclei moving through the stages of centrosome separation (i-ii), pronuclei migration and joining (iii), and pronuclear-complex rotation (iv). The bottom row shows the corresponding motor binding probabilities and the dynamic rearrangement of the terminator curves on the cell surface. The diffuse shadows in B-D are from graphics rendering with an external light source.

We assume Newton’s Third Law of equal and opposite forces governs the interactions between asters and pronuclei, assuming simple drag laws for their motion. In particular, the dynamics of each aster is governed by *η_a_****Ẋ****_a_*(*t*) = **F***_a_* = **F***_p_*_1_ + **F***_p_*_2_ + **F***_c_*, where *η_a_* is the drag coefficient of the aster, **F***_p_*_1_ and **F***_p_*_2_ are the pulling forces on the aster from pronuclear-bound motors, while **F***_c_* is that from cortical-bound motors. Similarly the dynamics of each pronucleus is governed by *η_p_****Ẋ****_p_*(*t*) = −**F**_1*p*_ – **F**_2*p*_, where *η_p_* is the drag coefficient of the pronucleus and **F**_1*p*_ and **F**_2*p*_ are the counter forces from the two asters on that pronucleus. Note that the drag-weighted force *η_a_*(***Ẋ****_a_*_1_ +***Ẋ****_a_*_2_) + *η_p_*(***Ẋ****_p_*_1_+***Ẋ****_p_*_2_) is balanced only by the total pulling force from the cortex. Hence, in the absence of cortical motors, the pronuclei/aster complex evolves as a force-free configuration. To calculate surface pulling forces in our model, we evolve coarse-grained surface fields for the binding probability of motors to microtubules from a particular aster.

How do motor binding probabilities evolve with space and time? We take the cortex as an example. The probability 𝒫*_ci_*(***Y***, *t*) of cortical motors at ***Y*** being bound to microtubules from aster *i* evolves with the local rates of microtubule binding to unoccupied motors and of microtubule detachment from the occupied motors:

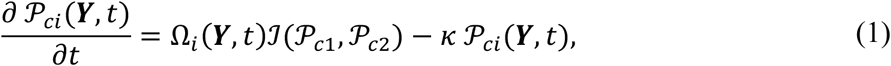

where the first term on the right-hand side gives the rate of binding, and the second the rate of unbinding with detachment rate *κ*. In particular, Ω*_i_*(***Y***, *t*) is the rate of microtubule impingement upon a motor domain centered at ***Y***, and ℐ(𝒫*_c_*_1_, 𝒫*_c_*_2_) represents the probability of a motor being unbound to either aster. This function ℐ captures the competition of asters for accessible motors.

Lastly, to account for line-of-sight interactions of asters with pronuclei and the cortex, we incorporate terminator curves, that divides motor-accessible from inaccessible regions, into the evolution equations of surface binding probability fields. Taking Eq. (1) for 𝒫*_ci_* as an example, if ***Y*** ∈ *S_c_* (the cortex) is, at time *t*, within a motor-inaccessible region for aster *i* we set Ω*_i_*(***Y***, *t*) = 0. There can be 12 distinct terminator curves, though not all can be realized simultaneously. These arise from each aster possibly producing terminator curves on each pronuclear surface *S_pj_*, and on the cell cortex *S_c_*, as well as shadowing between the pronuclei (terminator curves denoted, respectively, as *Γ_p_* and *Γ_c_* in Fig. 3B). The terminator curves on a pronucleus are a consequence of pronuclear convexity with respect to the aster positions and from shadowing by the other pronucleus. Those on the cortex arise solely from pronuclear shadowing. The terminator curves, both on the pronucleus and the cell cortex, are simply calculated by a cone whose apex is on the aster and which circumscribes the spherical pronucleus. These terminator curves evolve in time, constantly redefining the regions accessible to the astral microtubules. While there are now general computational methods for ray tracing with multiple surfaces which can shadow each other (*17*), our case is relatively simple given the spherical shape of the two pronuclei (Materials and Methods). Due to the mostly slow dynamics of the pronuclei and asters, we do not account for the finite speed of growth of the microtubule plus ends in determining terminator curves. Figure 3C shows how the terminator curves affect surface motor binding probabilities given an arrangement of asters and pronuclei.

Having the motor binding probabilities on the two pronuclei and the cell cortex, the total force on an aster at ***X****_a_*, for a given arrangement of terminator curves, is

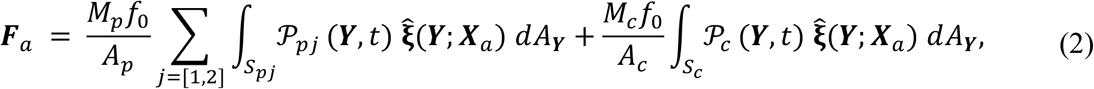

where the summation is over the two pronuclei, and *A_p_* and *A_c_* are the pronuclear and cellular surface areas, respectively.

We next test whether the GO-model, based on surface-anchored pulling acting through dynamically evolving terminator curves, is sufficient to reproduce pronuclear migration, meeting, centering, and rotation without introducing additional force-generating mechanisms. We perform and interpret simulations with asters and pronuclei initially arranged to mimic the intracellular geometry of early-stage *C. elegans* embryos: the male pronucleus is initially positioned near the posterior of the cell with the two asters caught in the narrow cleft between it and the posterior cortical surface. The female pronucleus is placed 40 μm away on the anterior side (Fig. 3D-i).

From their initial placements, the model asters attach to motors on both pronuclear and cortical surfaces, and are gradually pulled outwards, and away from each other. During this stage, there is little pronuclear motion. Indeed, the female pronucleus is motionless, as its surface is completely shadowed by the male pronucleus from the two asters. As they move out of the cleft between the pronucleus and the cell cortex, the asters gain access to more motors on the cortical surface that help pull them and the male pronucleus inwards, while also gaining access to motors on the female pronucleus, which initiates its migration towards the male pronucleus; see Figure 3D-ii, iii. The pronuclei subsequently meet, at which point we join them together as a single rigid body. As they join, the asters move into the groove between them, and the entire complex migrates toward the cell center and rotates to align with the anterior-posterior axis (Fig. 3D-iv, and movie S9).

All that was essentially done here was to model the geometry of microtubule impingement and pulling forces upon asters and pronuclei. What ensued was a dynamic qualitatively similar to that observed experimentally as the *C. elegans* embryo enters its first mitotic cell division.

### The GO-model explains the dynamics of asters and pronuclei

To determine if the GO-model accurately describes aster and pronuclear dynamics in *C. elegans*, we acquired 3D time-lapse movies from experiments spanning from initial aster separation to completion of pronuclear rotation (Fig. 4A and movie S10). By tracking the centrosomes and segmenting the pronuclear envelopes, we measured three dynamical processes: (1) aster separation, as measured by the angle formed by the two centrosomes relative to the center of the male pronucleus (Fig. 4B-ii); (2) pronuclear migration, measured by the distance between the centers of the male and female pronuclei (Fig. 4B-iii); and (3) the pronuclear-complex rotation, quantified by the orientation of the line connecting the two centrosomes relative to the cell’s anterior-posterior axis (Fig. S4). Experimentally, we found that the two asters initially separate at an angular speed of ∼ 0.4*π* rad/min, after which the female pronucleus migrates toward the male pronucleus at speeds around 20 µm/min. Following the meeting of the pronuclei, the asters reposition themselves into the groove between them, and the pronuclear complex moves toward the cell center while rotating with an angular speed of ∼ 0.1*π* rad/min. Once positioned at the cell center, the complex remains stable through spindle formation. Figure 4B compares the experimental measurements characterizing these dynamical processes with a simulation of the GO-model; this single simulation transitions from aster separation to pronuclear migration to centering and rotation, giving results that closely match the control experimental results (n=20) (Fig. 4B-ii, B-iii, and Fig. S4).

**Fig. 4.**
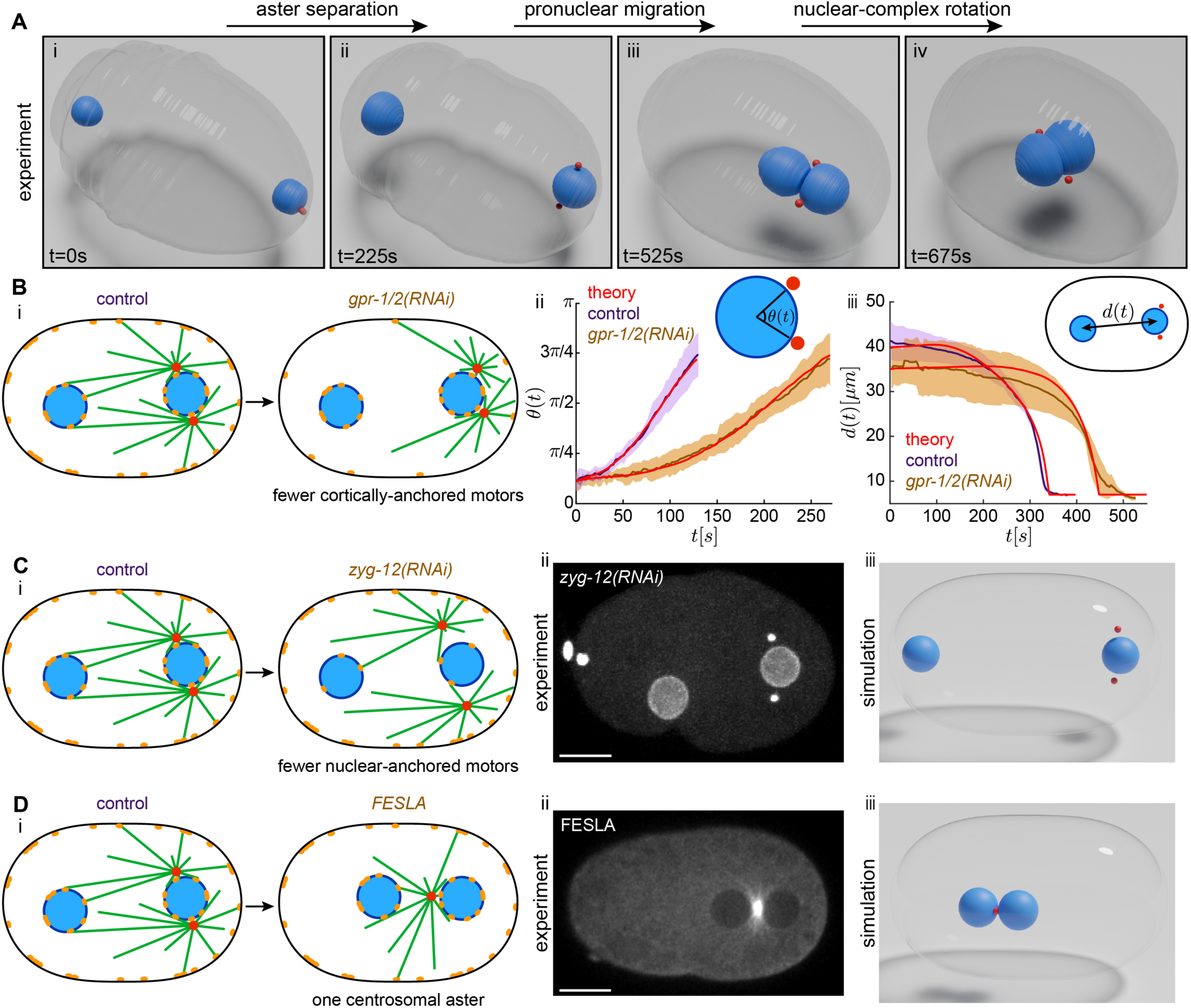
The GO-model explains aster and pronuclei dynamics in control and perturbed embryos. (**A**) Snapshots showing 3D motion of centrosomes (red) and pronuclei (blue) segmented from live imaging of an early *C. elegans* embryo. (**B**) (i) Schematic of *gpr-1/2(RNAi)*, where the density of cortically anchored motors is reduced relative to the control. (ii) Aster separation angle as a function of time for control embryos (purple), and *gpr-1/2(RNAi)* embryos (orange), and the simulated GO-model (red). The inset defines the separation angle, *θ*(*t*), measured relative to the center of the male pronucleus. For experiments, solid lines indicate the average; shaded regions denote the standard deviation (n=24). (iii) Separation distance of pronuclei during migration as a function of time for control embryos (purple), and *gpr-1/2(RNAi)* embryos (orange), and the GO-model simulation (red). The inset defines the inter-nuclei distance *d*(*t*) between the pronuclear centers. For experiments, solid lines indicate the average; shaded regions denote standard deviation. (**C**) (i) Schematic of *zyg-12(RNAi)*, where the density of pronuclear-anchored motors is reduced relative to control. (ii) A snapshot from an embryo dissected from a worm after four hours on *zyg-12(RNAi)* plate. Scale bar, 10 μm. (iii) Snapshot from a simulation of the GO-model with reduced pronuclear motor density. (**D**) (i) Schematic of FESLA experiment in which one centrosome is permanently ablated. (ii) A snapshot of an embryo after FESLA showing the remaining centrosome moving into the space between the two pronuclei. Scale bar, 10 μm. (iii) A snapshot from a simulation of the GO-model with one centrosome and two pronuclei, where the centrosome moves to the space in between the two pronuclei).

The quantitative agreement between the GO-model and experiment suggests that the model can be used to probe how force generation evolves within the cell as asters and pronuclei move and arrange themselves. In the GO-model, each terminator curve separates sets of motors engaged by a single aster (single source illumination), motors engaged by both asters (double source illumination), and disengaged motors (in total shadow) which exert no force upon the asters. In a singly illuminated region, motors only pull upon its illuminating aster and with the magnitude of force that depends upon the distance and orientation of region relative to the aster, as well as the assumed exponential length distribution of microtubules. In a doubly-illuminated region, motors are shared in the sense that a particular motor pulls upon one aster, or the other, but not both, and so the region’s total pulling potential is divided. In previous work, we showed that this competition for motors in a doubly-illuminated set could lead to an effective repulsion between the asters (*10, 16*). In our simulation, these sets dynamically appear, expand, shift, and disappear as asters and pronuclei rearrange themselves within the cell.

To understand how this dynamic geometry translates into a determination of aster and pronuclear motion, we first focus on the initial separating of the two model asters in the narrow gap between the male pronucleus and the cell cortex. Early on, their closeness causes the asters to bind to overlapping pools of motors on both the male pronuclear surface and the cell cortex. Their competition for these motors leads to a force imbalance, as sets where motors are bound to only one aster (singly illuminated) can give their undivided pulling force. This drives aster separation with each aster moving outwards towards a region of unshared motors. To further examine the role of competition in aster separation, we repeated the simulation with the same initial conditions and parameters but with only one aster. In this case, the single model aster moves very slowly, if at all (Fig. S5-B). We also simulated the system with reduced cortical motor densities, and thus, less competition, and observed a pronounced slowdown in aster separation (Fig. S5-C). These two GO-model simulations suggest that competition for cortical motors is central to timely aster separation.

To experimentally test the effect of motor competition on aster separation and pronuclear migration, we measured these dynamics in *gpr-1/2(RNAi)* embryos, where dynein anchoring to the cortex is impaired, and cortical pulling forces are weakened. Aster separation speeds were reduced by approximately twofold (Fig. 4B-ii, brown curve, movie S11) in these embryos - a result quantitatively reproduced in the GO-model by a ∼3-fold decrease in cortical motor density (Fig. 4B-ii, red line and brown). In the same simulation, slower aster separation delayed “illumination” and terminator curve formation on the female pronucleus, consequently slowing its migration (Fig. 4B-iii, red curve). That is, a reduction in only motor density was sufficient to quantitatively explain slower aster separation and delayed pronuclei migration. These findings align well with prior observations that simultaneously reducing both cortical and pronuclear motor densities diminishes the rate of aster separation even further (*8*).

We then used the GO-model to ask how the migration of the pronuclei is orchestrated. Recall that we used FESLA to repeatedly ablate the astral microtubules between the two pronuclei. This halted the posterior-directed migration of the female pronucleus and caused the asters to snap back to opposing positions on the male pronucleus (Fig. 2D-ii, red arrows). Within the GO-model, as the asters migrate to opposing sides of the male pronucleus, the female pronucleus also moves towards the male. At this onset time (to female migration), *t=t*^∗^, we performed a similar *in silico* “ablation” by setting the impingement rate of astral microtubules, Ω*_i_*(***Y***, *t*), from both asters to zero for *t* > *t*^∗^, on the female pronucleus and the anterior cortical surface. As in the experiment, due to the loss of pulling forces between the female and male pronuclei, this stops the migration of both pronuclei towards each other (Fig. S5-D). To specifically evaluate the influence of pronuclear-bound motors, we partially depleted *zyg-12*, a protein crucial for anchoring dynein to the pronuclear envelope (Fig. 4C-i). Extended depletion (> six hours) resulted in pronounced but highly variable phenotypes (movie S11). Thus, we limited depletion to four hours, and consistently observed asters positioned farther from the male pronuclear surface during their separation (Fig. 4C-ii and movie S11). Model simulations reproduced this phenotype when its pronuclear motor density was reduced ∼15-fold (Fig. 4C-iii). Lastly, we permanently removed one aster using FESLA. The remaining aster then repositioned itself along the axis connecting the two pronuclei (Fig. 4D-i, D-ii, and movie S12), in contrast to the off-axis arrangement observed with two asters (Fig. 1C). Simulations of the model in which one aster was removed again captured this experimentally induced single-aster behavior (Fig. 4D-iii).

## Discussion

To summarize, electron tomography, targeted laser ablation and other perturbations, and biophysical modeling all point to a self-organized mechanism for pronuclear migration, merger, centering, and rotation—central steps in reorganizing the zygote from fertilization to the first cell division. In this mechanism, the evolving geometry of pronuclei and asters is inseparable from the evolving geometry of the forces that move them: line-of-sight constraints gate where surface-bound force generators can engage, a geometric-optics analogue captured by dynamic terminator curves. This feedback loop connecting geometry, access, and force provides a unified physical explanation for reliable intracellular migration and positioning, an intricate process involving multiple organelles.

Understanding aster and nuclear positioning has spurred cross-disciplinary work in biology, physics, and applied mathematics (*18–22*). By casting pronuclear positioning in terms of a geometry–force feedback loop, our study provides a compact theoretical framework for integrating disparate observations across stages and perturbations. For example, in *C. elegans* the earliest displacement of the female pronucleus is driven by cytoplasmic flows generated by actomyosin-based cortical contractions (*23*). Although such flows are not directly modeled here, they can be incorporated naturally via the coarse-graining of fluid–structure interactions (*24, 25*), enabling future work that integrates cortical dynamics and cytoplasmic flows with aster-mediated organelle positioning.

Finally, our study invites an evolutionary question: how universal are such geometric feedback-loop mechanisms across species, where cell shapes and sizes can be highly diverse? Previous works has examined scaling of spindle elongation dynamics with cell size across a range of nematode species (*26, 27*) and across broader metazoan lineages (*28*). Using a stoichiometric pulling model for aster positioning, we showed that this interspecies diversity, at least among nematode species that are hundreds of millions years apart, can be explained by biophysical principles grounded in motor/astral-microtubule interactions and cell geometry (*10*). As in nematodes, the migration and union of female and male pronuclei after fertilization is a conserved evolutionary phenomenon across widely-separated species such as sea urchin (*29*), flies (*30*), and mouse and human (*31*). We anticipate that geometric feedback-loop mechanisms provides a common foundation for understanding these early events in animal development.

## Author contributions

Conceptualization: RF, DJN, TMR, MJS

Methodology: RF, YNY, DJN, TMR, MJS

Investigation: RF, GF, HYW, YNY, DJN, TMR, MJS

Visualization: RF, GF, JV

Funding acquisition: DJN, TMR, MJS

Writing – original draft: RF, MJS

Writing – review & editing: RF, GF, DJN, TMR, MJS

## Data and materials availability

All data are available in the main text or the supplementary materials.

## Supplementary Materials

### Materials and Methods

#### Maintenance of strains and fluorescence light microscopy

*C. elegans* strains were cultured at 24°C on nematode growth media (NGM) plates seeded with *Escherichia coli* OP50 as described (*32*). After thawing, the animals were propagated for approximately one week before imaging. Adult hermaphrodites were dissected in M9 buffer. Embryos were mounted on 4% (w/v) agarose pads between a slide and a coverslip. An eyelash was used to position selected embryos on the agarose pad. For live-cell microscopy, a spinning disk confocal microscope (Nikon TE2000, Yokugawa CSU-X1) equipped with 488 nm and 561 nm diode lasers, an electron-multiplying charge-coupled device (Hamamatsu, ImagEM Enhanced C9100-13), and either a 60× water-immersion objective (Nikon, CFI Plan Apo VC 60× WI, NA 1.2) or a 40× water-immersion objective (Nikon, CFI Plan Apo VC 40× WI, NA 1.25) was used. A home-developed LabVIEW program (LabVIEW, National Instruments) was used to control the parameters for imaging. For 3D tracking of centrosomes and pronuclei, we acquired 19 z-planes separated by 0.75 µm in 3-second intervals. Data for the nuclear-complex rotation (Fig. S2) was taken on a Nikon AX laser scanning confocal microscope with the NIS-Elements software using a 40×/1.2 silicon oil objective lens and pinhole set to 1 Airy unit.

#### High-pressure freezing, freeze substitution, and serial sectioning for electron microscopy

Single *C. elegans* embryos were ultra-rapidly frozen using an HPF Compact 03 high-pressure freezer (Engineering Office M. Wohlwend, Switzerland). For each freezing run, approximately five N2 adult hermaphrodites were dissected in a droplet of M9 buffer containing 20% (w/v) bovine serum albumin (Roth, Germany). An individual early embryo at the stage of pronuclear appearance was sucked into a cellulose capillary tube mounted into a pipette tip (*33*). The capillary containing the embryo was trimmed down to a final length of about 1-2 mm and sealed with two dull scalpels. The embryo within the capillary tube was then observed until it reached the desired developmental stage. Using fine tweezers, the capillary tube with the staged embryo was then transferred to a hexadecene (Merck, Germany) pre-wetted 100 µm-deep cavity of a type-A aluminum planchette (Wohlwend, article #241) filled with M9 buffer containing 20% (w/v) bovine serum albumin (Roth, Germany). The specimen holder was gently closed by placing the flat side of a type-B aluminum planchette (Wohlwend, article #242) on top of a type-A specimen holder. Such a ‘sandwich’ was then frozen under high pressure (∼2000 bar) with a cooling rate of ∼20,000°C/s (*33*).

Specimen holders were opened under liquid nitrogen and transferred to cryo-vials filled with anhydrous acetone containing 1% (w/v) osmium tetroxide (EMS, USA) and 0.1% (w/v) uranyl acetate (Polysciences, USA). Freeze substitution was performed in a Leica AFS2 (Leica Microsystems, Austria). Samples were kept at −90°C for one hour, then warmed up to −30°C with steps of 5°C/h, kept for 5h at −30°C, and warmed up again in steps of 5°C/h to 0°C. Afterwards, the samples were washed three times with pure anhydrous acetone and infiltrated with Epon/Araldite (EMS, USA) epoxy resin at increasing concentrations of resin (resin:acetone: 1:3, 1:1, 3:1, then pure resin) for 2h each step at room temperature. Samples were incubated with pure resin overnight and then for 4h. Then, samples were thin-layer embedded between two Teflon-coated glass slides and allowed to polymerize at 60°C for 48h (*34*). Polymerized samples were remounted on dummy blocks and semi-thick serial sections (300 nm) were cut using an ultramicrotome (EM UC6, Leica Microsystems, Austria). Ribbons of serial sections were collected on Formvar-coated copper slot grids, post-stained with 2% (w/v) uranyl acetate (EMS, USA) in 70% (v/v) methanol and then with 0.4% (w/v) lead citrate (Science Services, USA) in double-distilled water (*33*). In preparation for electron tomography, both sides of the samples were coated with 20 nm colloidal gold (BBI, UK), functioning as fiducial markers for tomographic reconstruction.

#### Electron tomography and 3D reconstruction

Serial sections containing early embryos were pre-inspected at low magnification (∼2900×) using a Zeiss EM906 transmission electron microscope (Zeiss, Germany) operated at 80 kV and equipped with a 2k×2k slow-scan CCD camera (Sharp eye, TRS, Germany). Serial sections containing regions of interest were mapped and then transferred to a transmission electron microscope (Tecnai F30, Thermo Fisher) operated at 300 kV and equipped with a 4k×4k CMOS camera (OneView, Gatan, USA). Using a dual-axis specimen holder (Type 2040, Fishione, USA), tilt series were recorded from −60° to +60° with 1° increments at a magnification of 4700× and a pixel size of 2.6 nm, applying the SerialEM software package (*35*). For dual-tilt electron tomography, the grids were rotated for 90° in the XY-plane and a second tilt series was acquired using identical microscope settings. The resulting tomographic A- and B-stacks of the same positions were reconstructed, combined, and flattened using IMOD (*36*).

#### Segmentation of microtubules and pronuclear membranes in electron tomograms

In each individual reconstruction all MTs longer than 100 nm were automatically segmented using the ZIB Amira (Zuse Institute Berlin, Germany) software package. After manual correction of the MT segmentation, the individual tomograms of each recorded embryo were stitched using the segmented MTs as initial landmarks to represent the whole microtubule network in the resulting 3D model (*37*). The pronuclear envelopes and the centrioles were manually segmented using the ZIB Amira software.

#### Femtosecond stereotactic laser ablation (FESLA)

The femtosecond laser pulses (∼70 fs pulse width and 800 nm center wavelength) from an ultrafast Ti:Sapphire laser (Mai Tai DeepSee, Spectra-Physics) were reduced from 80 MHz to 16 KHz by a pulse picker (Eclipse, KMLabs). The 16 KHz pulse train was directed into the spinning disk confocal microscope (Nikon TE2000, Yokugawa CSU-X1) and focused onto the sample by a high-numerical-aperture water-immersion objective (Nikon, CFI Plan Apo VC 60× WI, NA 1.2) for ablation. FESLA was performed through coordinated 3D sample movement (by Physik Instrumente P-545 PInano XYZ piezo-stage) and synchronized laser exposure (by Newport mechanical fast shutter). A home-developed LabVIEW program (LabVIEW, National Instruments) was used for image acquisition and FESLA execution.

#### Tracking centrosomes and segmentation of the pronuclei in live microscopy

We develop a MATLAB package to find the 3D position of the centrosomes and segment the pronuclei in live microscopy images. In short, to locate centrosomes’ positions at subpixel resolution, we performed Gaussian fitting. For one time point, the position of the centrosomes is manually determined. Then the software uses those positions as an initial guess for fitting a Gaussian function to the next frame. If the graphical user interface allows the user to manually correct and repeat this procedure if needed.

To segment the 3D shape of the pronuclei, we use active contour method (REFs). In short, we start with a 3D mesh with a shape of sphere in the center of the pronucleus as the initial guess. We then evolve the mesh based on an energy function that has a term associated as an internal pressure, a term associated with surface tension to keep the surface smooth, and a term that interacts with the image to find the edge of the pronucleus. The parameters are set such that the surface evolves while the mesh is inside the pronucleus and stop expanding when it reaches the surface of the pronucleus. The final shape of the mesh is then set by the minimum of the energy. In our software, we use BasicSnake_version2f written by D. Kroon from University of Twente.

Below is a snapshot of the graphical user interface.

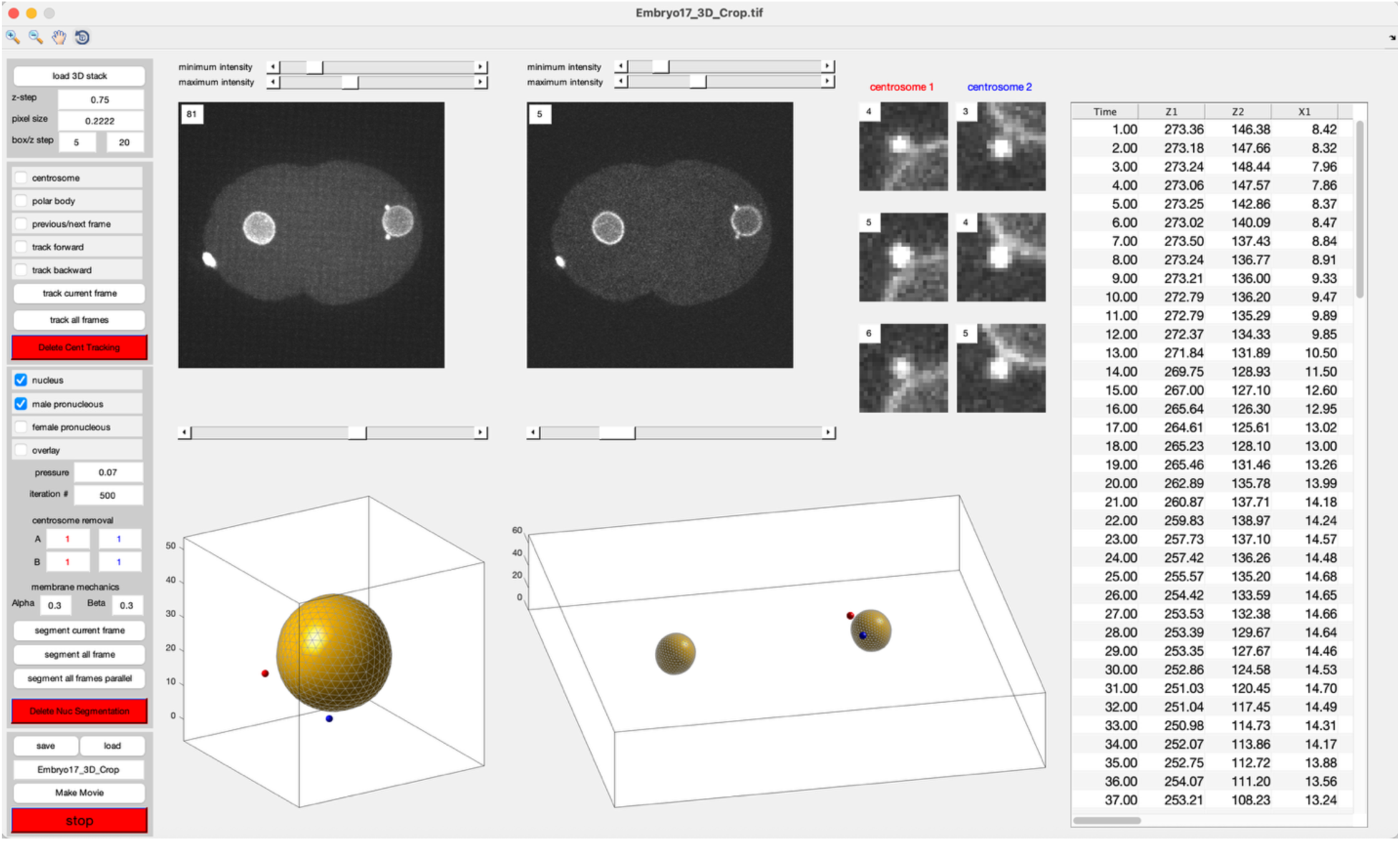

### Supplementary Text

#### Formulation of the geometric optics (GO) model

In the GO-model, we account for cell shape, two mobile centrosomal asters, and two mobile and spherical pronuclei. All surfaces (cell periphery and pronuclear surfaces) are loaded with dynein-type motors. The interaction of the microtubules with a motor is stoichiometric, constraining the motor to bind to only one microtubule at a time. We assumed that motors on the pronuclear and cell surfaces have the same biophysical properties. We first present the equations of motion for the asters interacting with both cortical and nuclear motors. We then present the equations of motion for the pronuclei, and finally, we formulate the computation of the terminator curves.

Consider two asters at positions 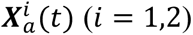, where microtubules nucleate from each aster at a rate *ṅ*, grow with a speed *V_g_*, and undergo catastrophe at a rate *λ*. The length distribution of the microtubules emanating from each aster is given by 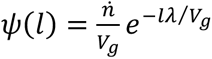. From those microtubules that reach an unbound motor at position **Y** (on the cell surface or pronuclear surfaces), only one microtubule binds to the motor and gets pulled upon. The force on this microtubule and the corresponding aster is given by 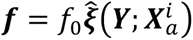, where *f*_o_ is the magnitude of the force from a bound motor and 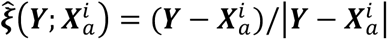 is a unit vector along the microtubule. The motion of the asters under forces from bound motors on the two pronuclear surfaces and the cell surface is governed by

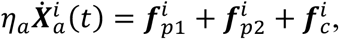

where *η_a_* is the drag coefficient for the aster, 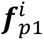 and 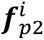 are the total pulling forces on the *i*th aster from the pronuclear motors, and 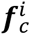 is the total pulling force from the motors on the cell periphery.

These forces are calculated from the surface fields, 𝒫*^i^*(**Y**, *t*), indicating the probability of binding of a microtubule from the *i*th aster to a motor at position **Y** at time *t*. The total force on the *i*th aster from the *j*th pronucleus, 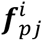, and from the cell periphery, 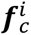, are given by

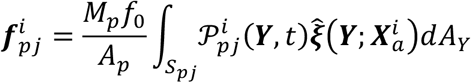

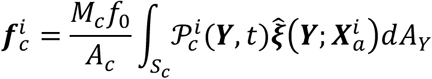

where *M_p_* and **M*_c_* are the total number of motors on the pronucleus and cell periphery, and *A_p_* and *A_c_* are the area of the pronucleus and the cell, respectively.

The time evolution of the probability fields 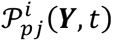 and are 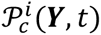 given by

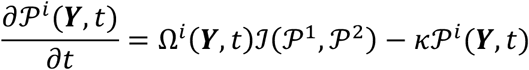

where Ω*^i^*(***Y***, *t*) is the impingement of microtubules from the *i*th aster on the point ***Y*** on the cell or the pronuclear surface, and *κ* is the motor unbinding rate. The function ℐ(𝒫^1^, 𝒫^2^) accounts for the stoichiometric binding of a motor to microtubules from the two asters and is given by

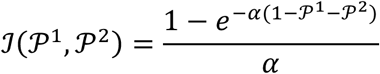

where 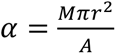 is the radius of the motor, and *A* is the total area of the corresponding surface (*A* =*A_p_*, for the pronuclei and *A* = *A_c_* for the cell surface). Similarly, *M* is the total number of motors on the corresponding surface (*M* = *M_p_* for the pronucleus and *M* = *M_c_* for the cell surface).

Next, we present the calculation of the microtubule impingement rate. For a point ***Y*** on the cell surface and at a distance *d* from the *i*th centrosome, the impingement rate 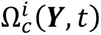 is

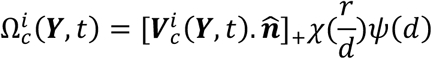

where 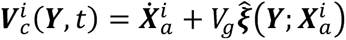 is the velocity of the microtubules’ plus ends relative to the cell surface and 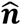 is the unit outward normal vector. The geometric factor 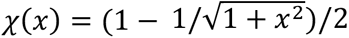 and the []_+_ sign ensure that the impingement rate always remains positive.

For a point ***Y*** on the *j*th pronucleus surface and at a distance *d* from the *i*th centrosome, the impingement rate 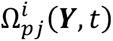 is

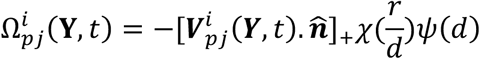

where 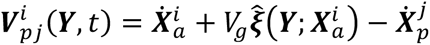 is the velocity of the microtubules’ plus ends relative to the *j*th pronucleus surface, with 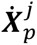 as the velocity of the *j*th pronucleus. The minus sign is due to microtubules impinging upon the pronucleus surface from outside.

Now, we describe the equation of motion of the two pronuclei. For the period that the two pronuclei are far from each other, we treat them as point objects with their equation of motion given by

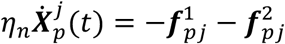

where *η_n_* is the drag coefficient for the pronucleus, and the right-hand side is the total counter-force from the two asters on the *j*th pronucleus. Once the two pronuclei get closer than the distance *∊*⁄2*R* ≪ 1 (*R* is the radius of the pronucleus), we consider the pronuclear complex as a rigid object with equations of motion

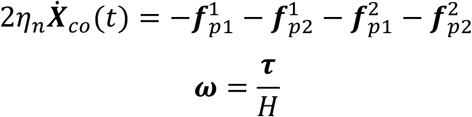

where 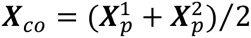 is the coordinate of the center of the pronuclear complex, and ***ω*** and ***τ*** are the angular velocity and total torque on the complex, respectively. *H* is the rotational drag on the pronuclear complex. The torque ***τ*** is given by

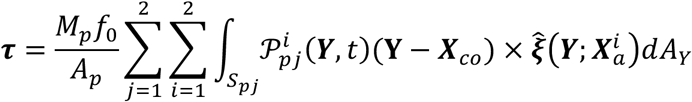

Following our tomography measurements, where we found that the number of microtubules increases by ∼20-fold during centrosome separation and pronuclear migration, we set the microtubule nucleation rate as

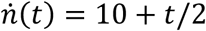

All the other parameters are time independent and are presented in Table 1.

**Table 1.**
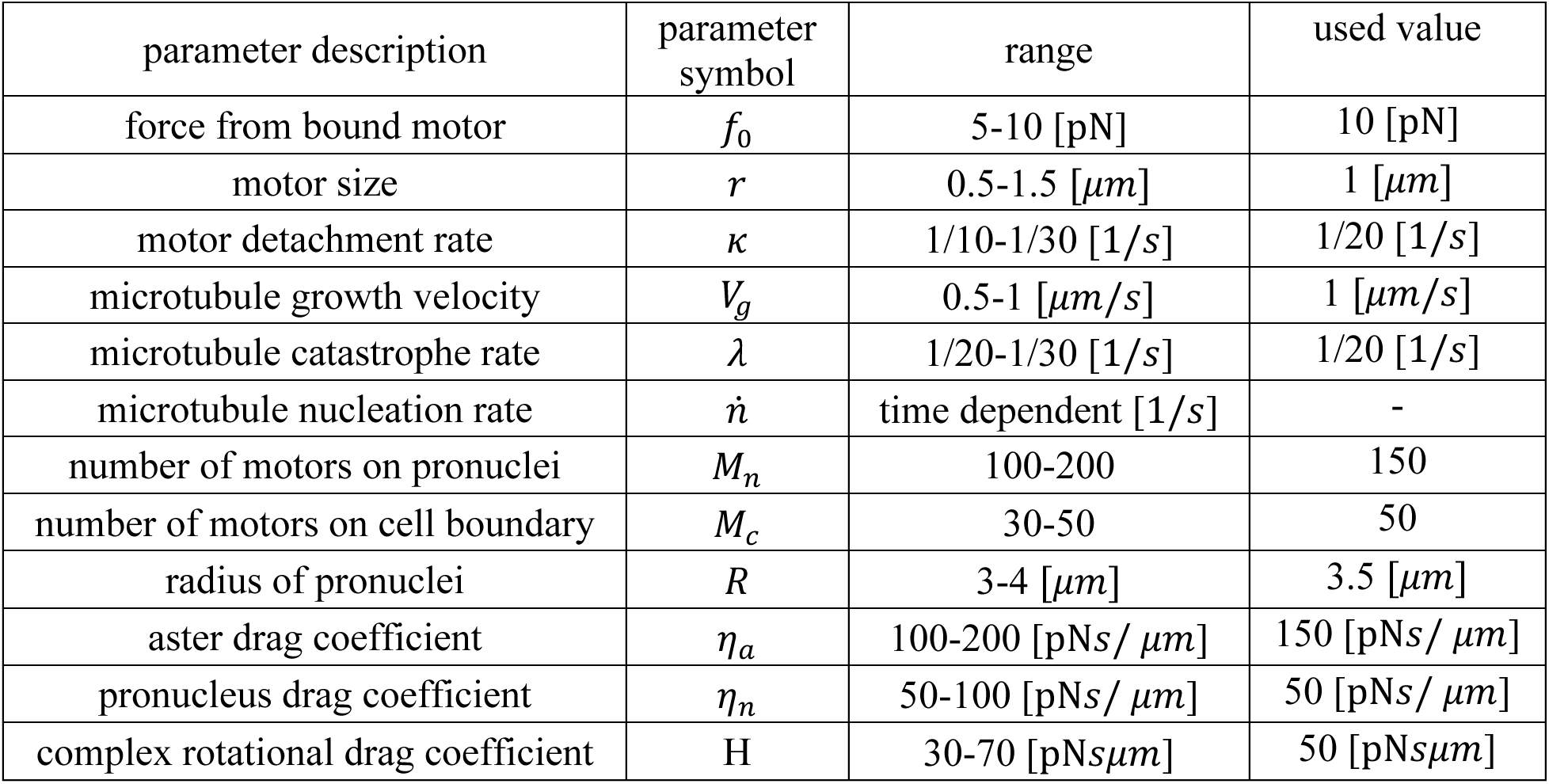
Parameters used for the simulation of geometric optics model.

#### Implementation of line-of-sight in the GO-model

Both the pronuclear surfaces and the cell cortex are represented as triangulated meshes, with each triangle defined by its center point and outward normal. To impose line-of-sight constraints, we use the concept of terminator curves, originating from ray-tracing, as the boundary between illuminated and shadowed regions of a convex surface relative to a point source. Each centrosome is treated as a point source, and the triangulated pronuclei and cell cortex are treated as convex surfaces. For a given centrosome position, the accessible region of a pronucleus is obtained by constructing the cone tangent to the pronucleus with its apex at the centrosome; the curve of tangency defines the terminator curve that separates illuminated (accessible) from shadowed (inaccessible) surface elements. Additional occlusion arises when the second pronucleus casts a shadow: triangles falling inside this shadow cone are also excluded. For the cell cortex, terminator curves are determined solely by occlusion from the pronuclei, again by projecting tangent cones from the centrosome. At each time step of the simulation, these terminator curves are recomputed geometrically, thereby partitioning triangulated surfaces into accessible and inaccessible regions along straight line-of-sight from the centrosomes. Below is the pseudo-code for the implementation of line-of-sight.

Input:

- Triangulated meshes of pronuclei and cell cortex (triangle centers + normals)
- Centrosome position(s)

For each centrosome:

For each pronucleus:

1. Construct tangent cone with apex at centrosome and tangent to pronucleus sphere
2. Terminator curve = intersection of cone with pronucleus surface
3. For each triangle:

a. Define direction vector

v = (triangle_center – centrosome_position) / ||triangle_center – centrosome_position||
b. Check orientation: accessible if dot(triangle_normal, v) ≤ 0
c. Check visibility: accessible only if triangle lies within illuminated region
4. Exclude triangles if they lie inside the shadow cone cast by the other pronucleus

For the cell cortex:

1. Project tangent cones from the centrosome through each pronucleus
2. Terminator curves = intersection of cones with the cortex
3. For each cortical triangle:

a. Define direction vector

v = (triangle_center – centrosome_position) / ||triangle_center – centrosome_position||
b. Mark triangle accessible if outside pronuclear shadow

Return:

- Accessible triangle indices on each surface for each centrosome

**Fig. S1.**
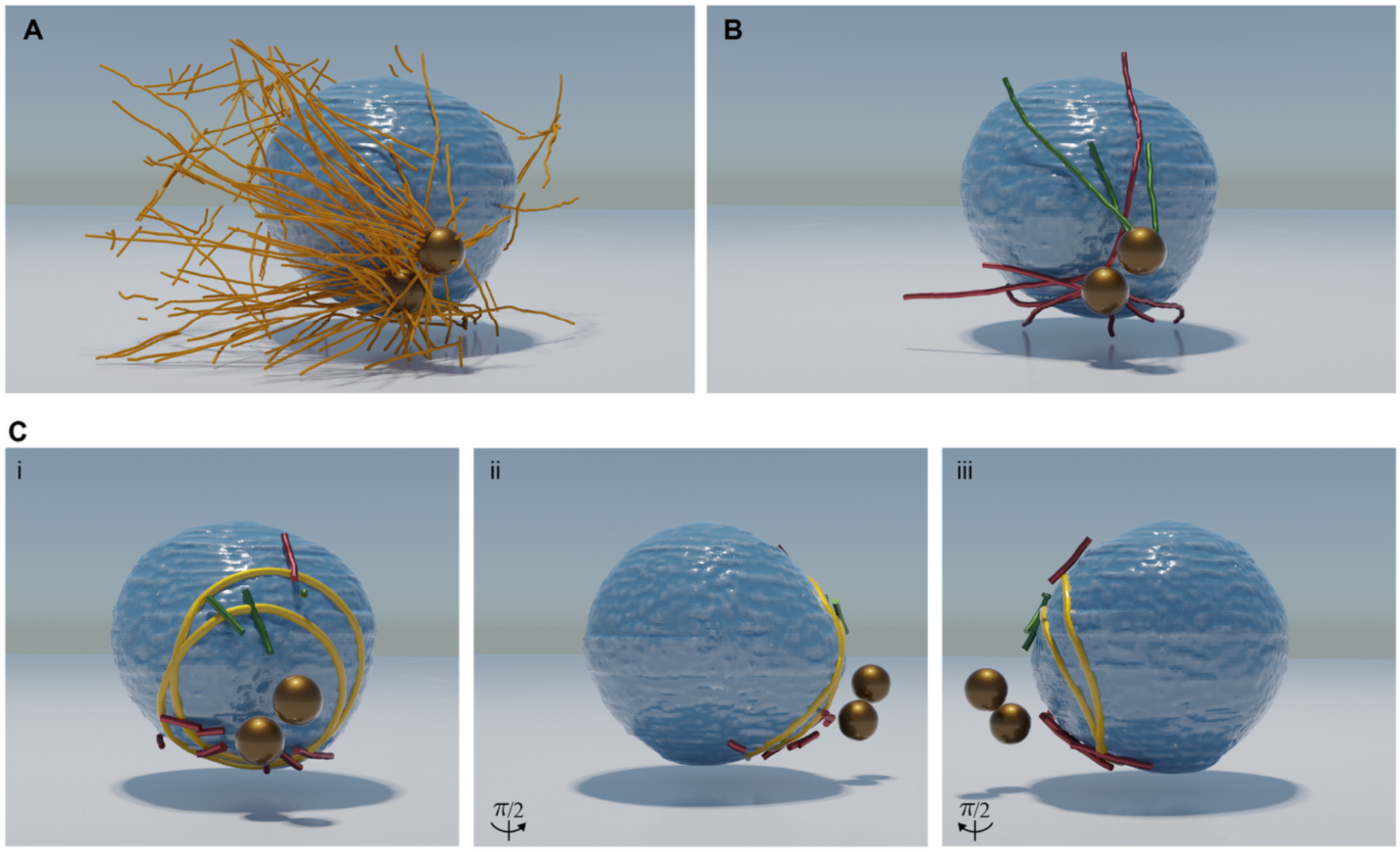
Microtubule organization during pronuclei migration. (**A**) 3D rendering of centrosomes (gold), microtubules (orange), and male pronucleus (blue) corresponding to Figure 1A. (**B**) Same rendering as in (A) highlighting only a subpopulation of microtubules that are 120 nm or closer to the pronucleus surface (microtubules from each aster are colored differently). (**C**) Same rendering as in (B) with microtubule segments that are 120 nm or closer to the pronuclear surface and the computed terminator curves (yellow) shown from different angles.

**Fig. S2.**
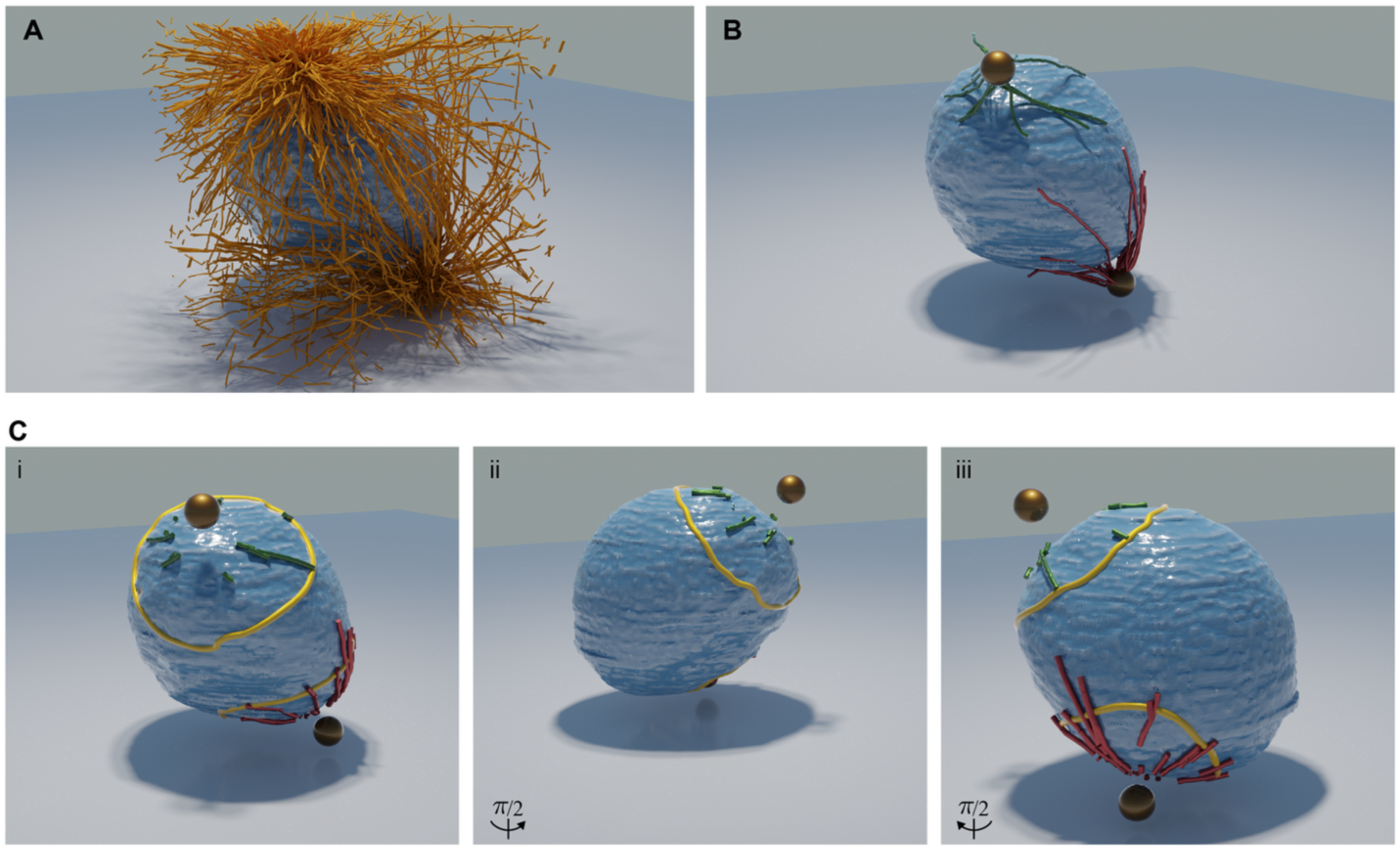
Microtubule organization during pronuclei migration. (**A**) 3D rendering of centrosomes (gold), microtubules (orange), and male pronucleus (blue) corresponding to Figure 1C. (**B**) Same rendering as in (A) highlighting only a subpopulation of microtubules that are 120 nm or closer to the pronucleus surface (microtubules from each aster are colored differently). (**C**) Same rendering as in (B) with microtubule segments that are 120 nm or closer to the pronuclear surface and the computed terminator curves (yellow) shown from different angles.

**Fig. S3.**
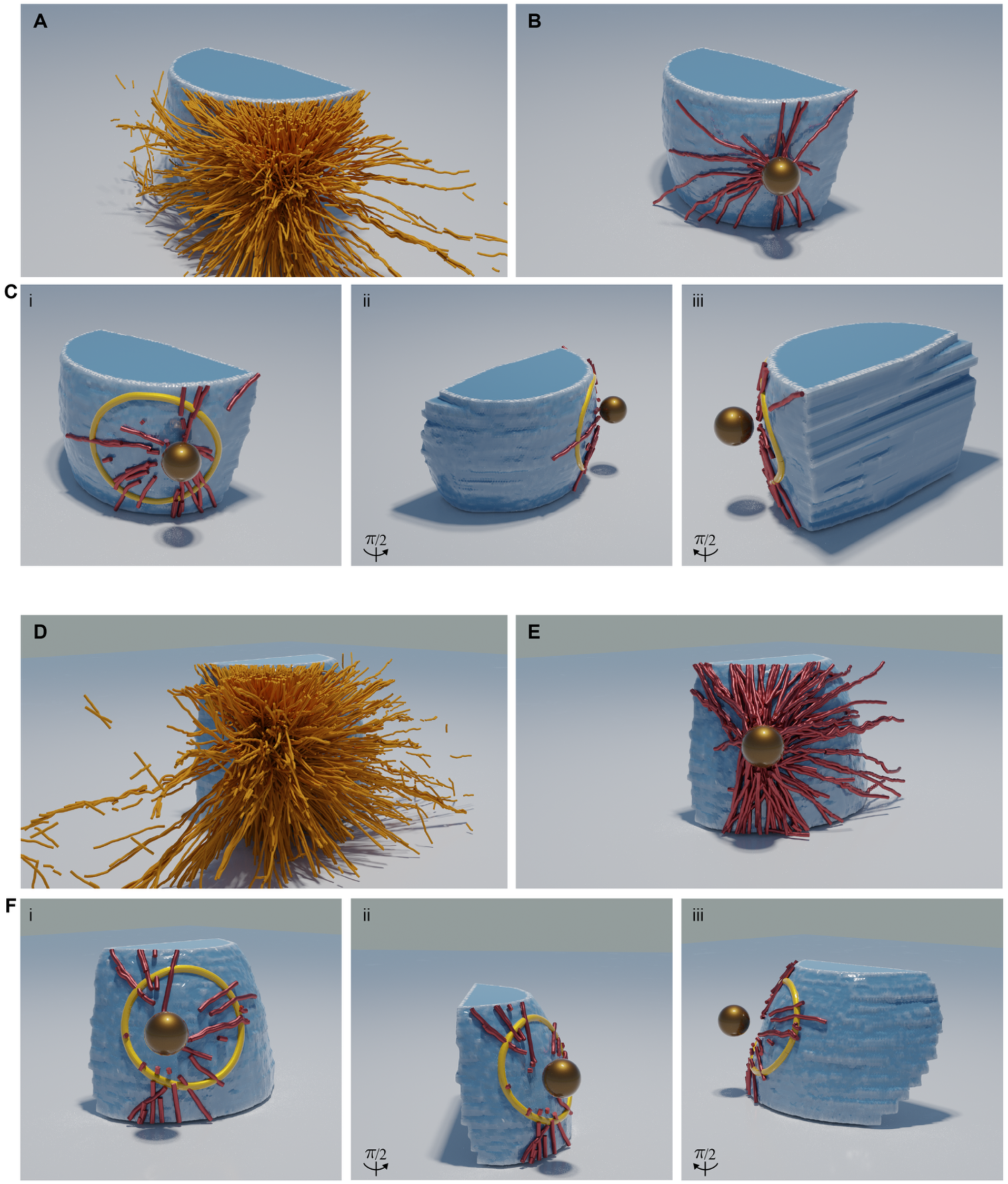
Microtubule organization during pronuclei migration. (**A, D**) 3D rendering of centrosomes (gold), microtubules (orange), and male pronucleus (blue) corresponding to Figure 1D. (**B, E**) Same rendering as in (A, D) highlighting only a subpopulation of microtubules that are 120 nm or closer to the pronucleus surface. (**C, F**) Same rendering as in (B, E) with microtubule segments that are 120 nm or closer to the pronuclear surface and the computed terminator curves (yellow) shown from different angles.

**Fig. S4.**
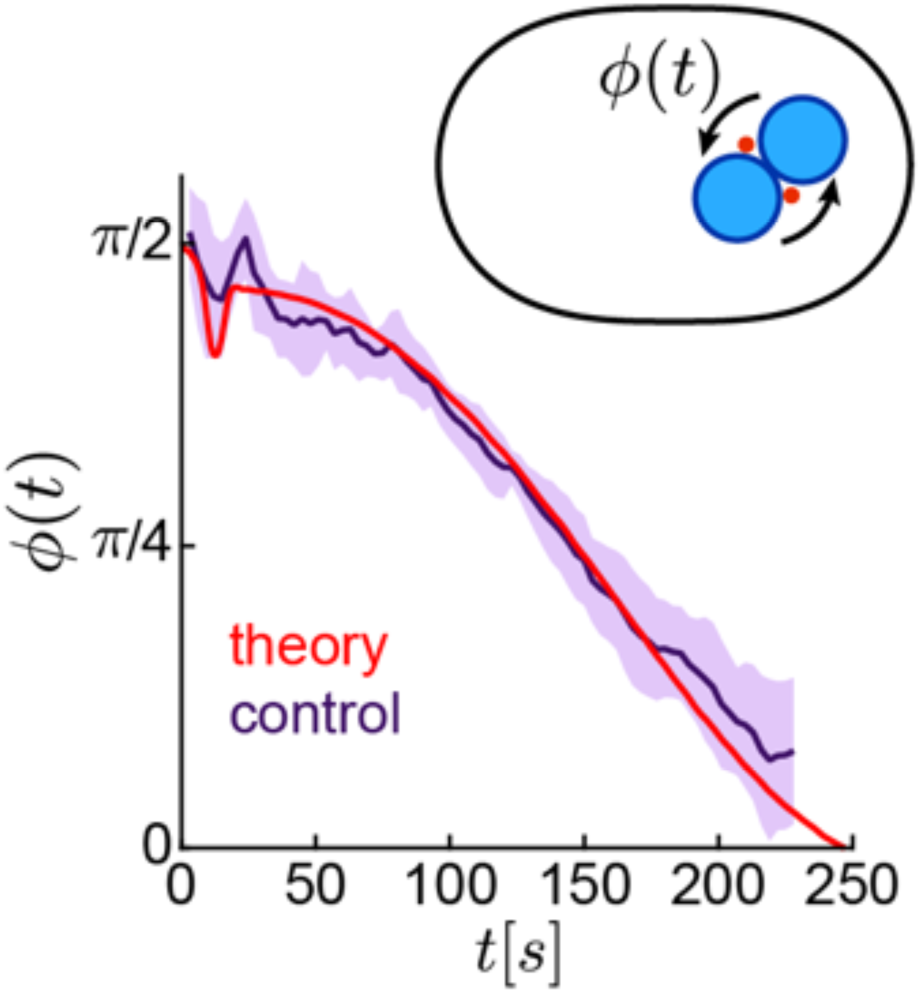
Geometric optics model explains nuclear-complex rotation. Angle of the nuclear-complex, measured as the orientation of the line connecting the two asters relative to the cell’s anterior-posterior axis, as a function of time. Theory is shown in red, and experiment for control embryos is in purple. For the experiment, solid lines indicate the average; shaded regions denote the standard deviation. The schematic shows the nuclear-complex angle.

**Fig. S5.**
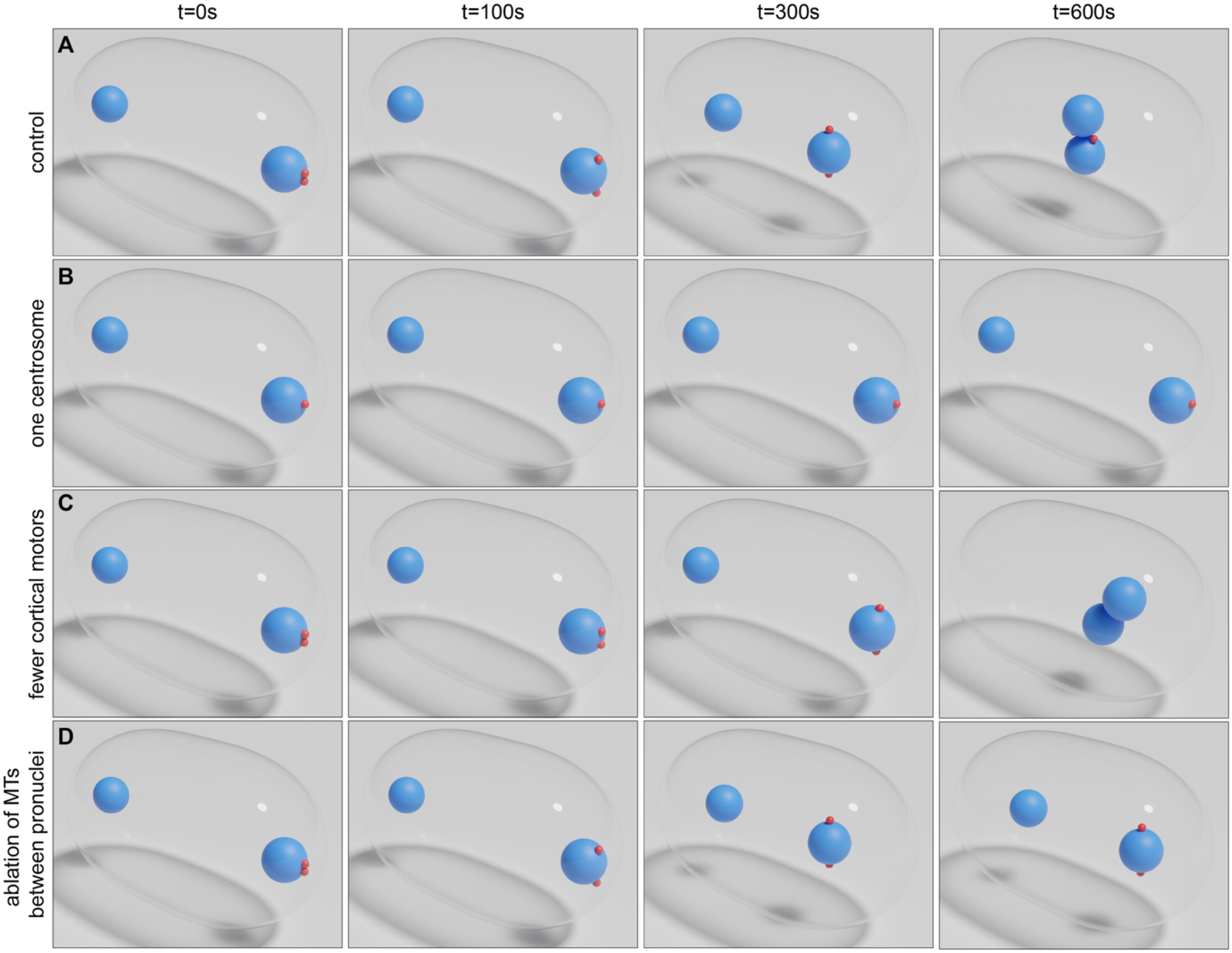
Simulation of geometric optics model for various conditions: (**A**) Snapshots from a simulation for control (see Table 1 for model parameters), (**B**) with the same initial conditions and model parameters as in (A) except with one aster, (**C**) with the same initial conditions and model parameter as in (A) except with 3-fold fewer cortical motors, and (**D**) with the same initial conditions and model parameter as in (A) except after the pronuclei reach the distance of ∼**25*μ****m* (at ***t*** = ***t***^∗^), we set the impingement rate to zero for all surfaces on the left side (*Ω* (***t*** > ***t***^∗^, ***x*** < **0**) = **0**).

**Movie S1.**

3D rendering of centrosomes (red), microtubules (orange), and the male pronucleus (blue) from electron tomography data shown in Figure 1A.

**Movie S2.**

3D rendering of centrosomes (red), microtubules (orange), and the male pronucleus (blue) from electron tomography data shown in Figure 1B.

**Movie S3.**

3D rendering of centrosomes (red), microtubules (orange), and the male pronucleus (blue) from electron tomography data shown in Figure 1C.

**Movie S4.**

3D rendering of centrosomes (red), microtubules (orange), and the male pronucleus (blue) from electron tomography data shown in Figure 1D.

**Movie S5.**

Severing of microtubules between the centrosome and the male pronucleus shown in Figure 2A.

**Movie S6.**

Severing of microtubules between the two asters shown in Figure 2C.

**Movie S7.**

Repeated severing of microtubules between the two pronuclei when they were ∼25 μm away from each other shown in Figure 2D.

**Movie S8.**

Severing of astral microtubules during the rotation of the pronuclear complex as shown in Figure 2E.

**Movie S9.**

Simulation of the geometric optics model for parameter values shown in Table 1. The left panel shows the centrosomal asters (red), pronuclei (blue), and the cell periphery (gray). The middle panel shows the motor binding probability on the cell periphery in the same simulation. The right panel shows the motor binding probability on the pronuclei in the same simulation.

**Movie S10.**

3D confocal microscopy (top panel) and tracking of centrosomes (red and blue) and pronuclei in a control *C. elegans* embryo.

**Movie S11.**

From left to right, live microscopy of centrosomes and pronuclei in control embryo, *gpr-1/2(RNAi)*, and *zyg-12(RNAi)*.

**Movie S12.**

FESLA experiment showing tub::GFP, where one centrosome is permanently removed.

